# Classification of down-core foraminifera image sets using convolutional neural networks

**DOI:** 10.1101/840926

**Authors:** Ross Marchant, Martin Tetard, Adnya Pratiwi, Thibault de Garidel-Thoron

**Affiliations:** Aix-Marseille Université, CNRS, IRD, Coll. De France, INRA, CEREGE, Technopôle de l’Arbois-Méditerranée, Aix-en-Provence, 13545, France

## Abstract

Manual identification of foraminifera species or morphotypes under stereoscopic microscopes is time-consuming for the taxonomist, and a long-time goal has been automating this process to improve efficiency and repeatability. Recent advances in computation hardware have seen deep convolutional neural networks emerge as the state-of-the-art technique for image-based automated classification. Here, we describe a method for classifying large down-core foraminifera image set using convolutional neural networks. Construction of the classifier is demonstrated on the publically available *endless forams* image set with an best accuracy of approximately 90%. A complete down-core analysis is performed for benthic species in the Holocene period for core MD02-2518 from the North Eastern Pacific, and the relative abundances compare favourably with manual counting, showing the same signal dynamics. Using our workflow opens the way to automated paleo-reconstruction based on computer image analysis, and can be employed using our labelling and classification software *ParticleTrieur*.

## I. INTRODUCTION

Foraminifera are cosmopolitan unicellular marine protists, very abundant in the ocean on both benthic and free-floating environments. They secrete unique carbonate shells, mostly on the submillimetric scale, that accumulate on the ocean floor, building kilometres of carbonate oozes. Sediment cores provide a record of foraminifera species composition and abundance over time. The presence of a species can be used to date marine sediments for biostratigraphy. The relative and absolute abundances of different species, along with their morphometric characteristics and geochemical composition, have been used for decades as proxies for reconstructing past climate conditions, such as heat content, oxygen concentration, and salinity of oceans (e.g. Kucera et al., 2012).

The processes required for acquiring these data necessitate the identification, and sometimes picking, of target species or morphotypes. However, this is often a time consuming manual process that needs to be performed by experts and requires advanced training. Typically, a sediment sample containing thousands of particles is placed under a microscope, through which a researcher manually identifies, counts and eventually picks the specimens of interest, usually at the species level. It can take many months or more to pick even a single species for a high-resolution geochemical analysis of a sedimentary record.

### A. Automated Identification

Robust, automatic identification of foraminifera and other micro-organisms such as coccolithophorids has thus been a subject of research over the last few decades (e.g., Liu et al., 1994; Culverhouse et al., 1996; Beaufort and Dollfus, 2004). The goal is to speed-up the identification process to reduce the time and cost of high-resolution studies, and improve reproducibility of classification, which can vary among researchers and is affected by experience level (Fenton et al., 2018). Shells of planktonic foraminifera retrieved from sediments have the peculiarity of being whitish and non-transparent, in contrast to living specimens. The specificity of the calcite in dead shells has the advantage of high-contrast on black backgrounds, making them ideal for optical imaging, yet some morphological features (internal or opposite to the field of image acquisition) cannot be seen due to this opacity.

Many approaches to the automatic classification of marine microfossils have been investigated. Morphological features obtained from image processing have been combined with rule-based (Yu et al., 1996), statistical (Culverhouse et al., 1996) or artificial neural network (ANN) classifiers (Simpson et al., 1992; Culverhouse et al., 1996, 2003; Hibbett, 2009; Schulze et al., 2013); images are directly input into systems such as the fat neural network used in the SYRACO (Dollfus and Beaufort, 1999; Beaufort and Dollfus, 2004) and the convolutional neural network (CNN) used in COGNIS (Bollmann et al., 2005); or both images and morphology combined (Barbarin, 2014). Of these methods, neural networks have shown superior performance to other statistical methods (Culverhouse et al., 1996). However, early attempts consisted of shallow CNNs with few convolutional layers that were time consuming to train, e.g. 30 hours for COGNIS on a 2000 image dataset (Bollmann et al., 2005) preventing an in depth-analysis of the robustness of those algorithms.

### B. Deep CNNs

Recent developments in computing power have reduced computation time of CNNs (Schmidhuber, 2014). At the same time, problems such as over-fitting (Hinton et al., 2012), where a CNN gives good accuracy on training images but not when applied to new unseen images, and vanishing gradients (He et al., 2015), where deep networks with many layers do not converge to a solution during training, have been addressed. This progress allows the construction of deeper networks (more layers) using larger images (e.g., He et al., 2015, 2016; Zagoruyko and Komodakis, 2016), and since 2012, the performance of deep CNNs on common evaluation datasets has surpassed engineered features (e.g. morphology) (Krizhevsky et al., 2012) and is approaching human performance (Russakovsky et al., 2015). Some popular networks include VGG (Simonyan and Zisserman, 2014), Inception (Szegedy et al., 2015b,a), ResNet (He et al., 2015, 2016; Zagoruyko and Komodakis, 2016; Xie et al., 2016) and DenseNet (Huang et al., 2016).

As a consequence, much research into using deep CNNs to automate image processing tasks in other fields is being performed. In the foraminifera domain, one current approach is using transfer learning with pre-trained ResNet50 and VGG networks to classify foraminifera images coloured according to 3D cues from 16-way lighting (Zhong et al., 2018; Mitra et al., 2019). Hsiang et al. constructed a large planktonic foraminifera image set, *Endless Forams*, using multiple expert input, and then applied transfer learning using the VGG network to compare CNN-based classification with humans (Hsiang et al., 2019).

At CEREGE, we have also been developing deep CNN classification systems for use in our microfossil sorting machine, MISO. The application is two-fold, firstly we wish to identify images so that the machine can separate the particle into different species or morphotypes for further analysis. Secondly, we want to classify images from large down-core datasets to perform species or morphotype counts and abundance calculations.

In this study, we detail our method for automated classification foraminifera images, with application to down-core images sets. The method is applicable to other single particle classification tasks. It consists of four steps:

- Acquisition of images.
- Curation of a training image set.
- Pre-processing the images
- Selection, training and evalution of a CNN
- Application of the CNN to classify a larger down-core image set.

In sections II to VII we explain the method, Section VIII gives a demonstration on the Endless Forams image set, and Section IX shows the results of analysis of benthic foraminifera for core MD02-2519.

## II. DOWN-CORE IMAGE ACQUISITION

The first step in the down-core analysis is to acquire images. The samples to be analysed are sieved to the desired size range, e.g. 150 *μ*m to 1 mm, and then split to approximately 3000 particles each. Each sample is either spread onto a micropaleontological and imaged with an automated microscope and stage, or using the microfossil sorting and imaging machine (MISO) at CEREGE.

For the imaging system, in both cases we use a 4× magnification telecentric lens (VS-TCH4) projecting onto an image sensor with 3.45 *μ*m wide square pixels (Basler acA2440-um) and illuminated with a white ring light at 30 degrees illumination angle (VL-LR2550W). This gives images with approximately 1159.4 pixels per millimeter resolution. The same camera exposure, gain and white balance are used for all images, typically 3000 ms, 0 db and 1.8 red : 1.0 green : 1.4 blue ratio, respectively.

The depth of field of our telecentric lens is approximately 90 *μ*m, and not enough to capture most foraminifera in our size range entirely in focus. Therefore fusing of multiple images at different focus depths (Z stack) is employed. Either the *Helicon* commercial image stacking software or our own custom algorithm is used to fuse the image into single full focus image (Figure 1). A separation of 70 *mu*m is used for the stack.

**Fig. 1.**
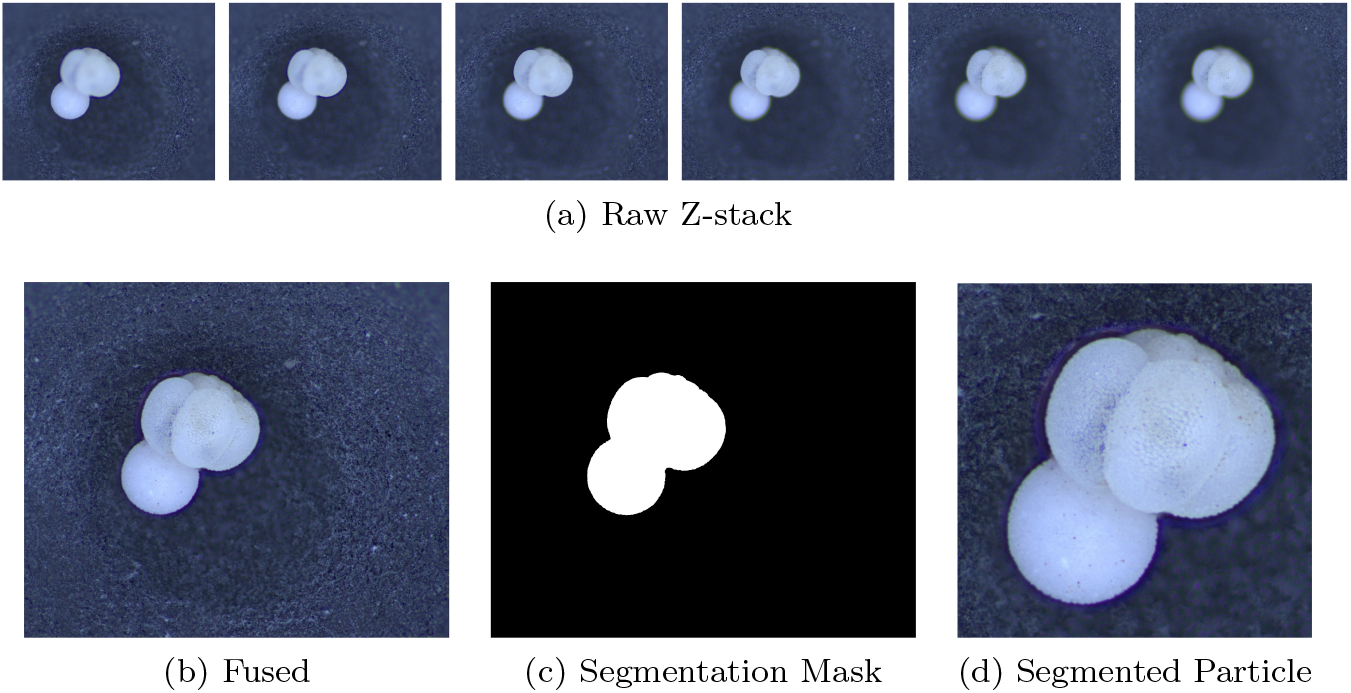
The image acquisition process. Raw images form a Z-stack (a) which is fused into an in-focus image (b), from which a segmentation mask is calculated (c) and the foraminifera particle is cropped (d).

Each foraminifera particle is then cropped into an individual image. Since foraminifera are generally bright white particles, a mask is found using binary segmentation of the image intensity using a fixed background threshold or a dynamic background model (MISO) (Figure 1c). The mask is smoothed using morphological opening and then separated into candidate regions of connected pixels. Regions that are too small or too locally concave, and thus not representative of a foraminifera shape, are removed. The remaining candidate regions are thus considered to represent particles.

The centre of mass (CoM), and the maximum radius between the CoM and the perimeter are calculated. Each particle is then segmented by cropping a square image centred at the CoM and with side length approximately 2.2 times the maximum radius (Figure 1d). This ensures the particles appear at roughly the same relative size in the images, with enough of a buffer to clearly realise the particle edge feature, as well as enable rotation of the particle within the image without going out-of-bounds.

Note that if using images from other sources, e.g. the *Endless Forams* database (Hsiang et al., 2019), any extra regions with non-photographed information, such as added white regions with metadata text, are removed.

## III. TRAINING SET CURATION

CNNs are a supervised learning method whereby the netwok must be trained to correctly identify images into various classes. A set of images, the training set, is created by choosing a number of example images for each of the desired classes. Typically, this involves a human expert familiar with the domain, who manually labels images with the correct class. The CNN is trained by feeding in batches of these images and calculating the difference between their predicted labels from the CNN and the true labels. The error is back-propagated through the network, updating the layer weights and other parameters to better discriminate the classes. With each subsequent batch, the predicted labels converge towards the true label values, until eventually there is no longer any improvement and training is considered complete.

Proper training set construction is therefore crucial for creating a good classifier. In particular, we draw attention to three main considerations:

1. CNNs learn to discriminate classes bases on the image features present in the training set. If the CNN is then used to classify other images from a particular class which do not contain these features, accuracy may be poor. For example, if only umbilical views of *N. dutertrei* are in the training set, dorsal views may not be classified correctly.
2. The CNN will learn any features that discriminate classes, even if they do not belong to the foraminifera. For example, if images of particular class are taken on a micropaleontological tray with rough background, while those for the other classes are taken on a tray with a smooth background, the CNN may learn that rather than the foraminifera shape, a rough image background is a distinguishing feature of the class, and then have difficulty classifying the same class on a smooth background, or other classes on a rough background.
3. The final layer of multi-class (each image has one label) CNN has dimension equal to the number of classes and typically uses a SoftMax activation function. That is, the output of the CNN is a value for each class, where the largest value corresponds to the predicted class, and that *all the values add up to 1.0*. Because of the later, if the CNN is used to classify images whose class is not in the training set, it can still be classified with high “probability” as belonging to one of the classes, especially if they share a distinguishing feature. Thus it is difficult to detect out-of-class images using the CNN output. This is especially noticeable when fragments and other non-foraminfera particles are present a sample but not in the training set.

### A. Variations

The training set should attempt to contain all the classes that we expect to encounter in the images to be classified. Furthermore, it should cover the intra-class variations that may be present, such as variation in particle appearance:

- Within-species morphological variability.
- Changes in preservation.
- Damage and fragmentation.

variations in the pose of the particle in the images:

- Aspect, for example umbilical, dorsal or lateral view.
- Rotation in the 2D image plane for a particular aspect.
- Position of the particle in the image.
- Size of the particle in the image.

and variations within the imaging system:

- Brightness, contrast, and colour shifts due to camera parameters, lighting brightness, colour and angle, and objective distortion or non-uniformity across the field of vision.
- Resolution and detail of the images.
- Other objects or background details in the background, e.g. tray surface.

Creating an image set that covers all permutations of these parameters would be time consuming. To ease the process we can attempt to remove different variances by pre-processing of the images, such as by normalising brightness and contrast, 2D orientation, particle size in the image, etc. Using the same camera, objective and imaging system for both creating the training set images as well as the classified images will also reduce variance.

Complementary to this, we can also add variance into the training set in a controlled and complete manner, by augmenting the initial training set with new images created by applying different transformations to the originals, such as gain (brightness), gamma (contrast) and particularly rotation. This makes the CNN more robust to these variations (e.g., Simard et al., 2003).

In general we prefer augmentation to pre-processing where possible, as pre-processing must be applied to images both before training and before classification, whereas augmentation is not needed for classification. This makes the classifier simpler to use and easier to share with other users, as they do not have to implement many pre-processing steps.

We use pre-processing to regularise the position and size of the foraminifera in the image, and fixed imaging parameters to reduce variance in brightness and colour. Augmentation is used primarily for robustness to 2D rotation, as well to brightness, contrast, size.

Other variations such as the within-class particle appearance and the 3D aspect (dorsal / umbilical / lateral view) can only be covered by the training set image selection.

### B. Labelling

With these caveats in mind, rather than trying to create a single universal foraminifera classifier, we create classifiers (and thus training sets), on a per-core or per-site basis. The idea being to ensure that the CNN is trained on the species or morphotypes that are specific to that core, and use images taken with the same acquisition system and camera parameters.

As such, a training set is chosen by selecting a images from a few representative samples, or taking a random subset from the larger down-core image set. Images are labelled with the aide of the *ParticleTrieur* software developed at CEREGE.

To begin with, images are labelled and new classes created as different species or morphotypes are identified. As labelling progresses, the number of images in each class is monitored, and low count classed are checked to make sure that the spectrum of morphological variability is covered. If there are not enough images in a particular class, more are added to the training set.

As a general rule, we aim for at least 200 images in each class, covering all the typical aspects (dorsal, lateral etc), although it can be difficult to find enough images for some rare classes. Once labelled, the images are exported in JPEG format into directories, one for each class. These form the training image set.

## IV. CNN TOPOLOGY

The completed training set is now used to train a CNN classifier. Ideally we want a classifier that:

- Gives high accuracy across all the classes to ensure good performance when applied to the larger down-core set.
- Is relatively quick to train so that any parameter tuning can be performed.
- Has a fast inference (classification of new images) time so that classifying the larger down-core image set is not too time consuming.

These objectives are realised by a combination of choice of CNN topology, training method, pre-processing and image augmentation.

We considered a variety of existing CNN designs when deciding which topology to use, as well as two custom topologies we have designed ourselves. We also consider employing transfer learning using pre-trained versions of some of the exisitng toplogies.

### A. Existing Topologies

Among the many CNN topologies proposed in the literature, some of the more popular systems include VGG, Inception, ResNet and DenseNet among others. These networks have been applied to large-scale problems such as classifying the ILSVRC image classification subset of ImageNet (**?**), which consists of 1.2 million images in 1000 non-overlapping classes of colour images.

We use the Keras open source library to create and train the CNNs. It is included with Tensorflow, a popular framework for machine learning developed by Google. Specifically, Keras provides versions of the VGG, ResNet, Inception, Densnet, MobileNet and NASNet networks, along with weights for these networks pre-trained on the ImageNet classification data base. We also another library, which provides a convenient interface to the Keras network and adds some smaller sized (18 and 34 layers) implementations of ResNet, as well as their squeeze excitation versions.

**Fig. 2.**
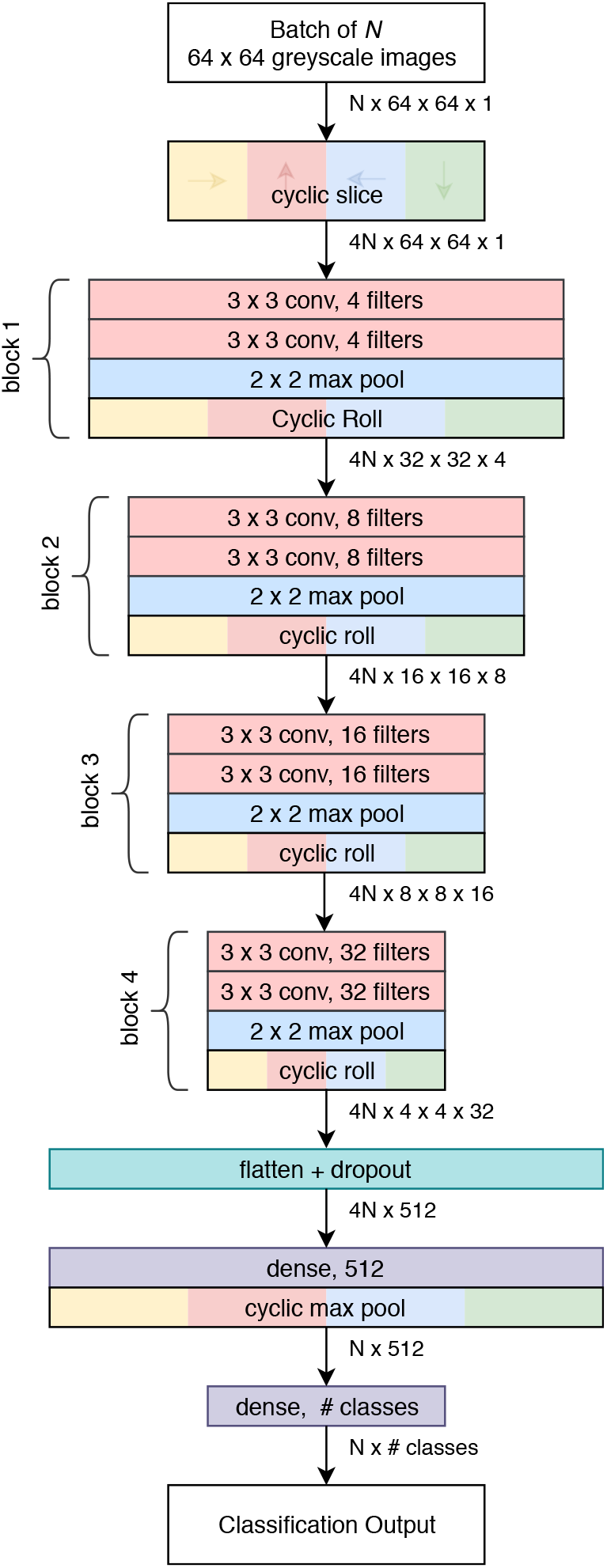
Topology of the *Base-Cyclic* CNN for an input image size of 64 × 64 pixels and four filters in the first block.

The available configurations of these CNNs have been developed with the ILSVRC ImageNet 2012 classification problem in mind (1.2 million colour images in 1000 non-overlapping classes). The size of the networks range from 3.5 million to 144 million parameters, with default image size ranges from 224 × 224 and 311 × 311 pixels in size. They have convolutional layers with many filters, presumably to have the capacity to handle large image sets like ImageNet.

### B. Custom Topologies

The aforementioned existing CNNs are designed for a large size input images and training sets with hundreds of classes. However, for foraminifera images can sometimes be quite small, in the case of large field microscopy, and to date our foraminifera classification tasks have consisted of fewer than 40 classes and fewer than 30000 images. As a consequence, we have also designed two new compact CNN topologies that can adapt to different input sizes.

Or first design, *Base-Cyclic*, uses convolutional units consisting of a 3×3 convolutional layer, followed by batch normalisation layer and then rectified linear activate (ReLU) [ref]. The convolutional layers are initialised using He’s method [ref] as this was found to improve training convergence. Two convolutional units are combined into a block, with a 2×2 max pooling layer at the end. A network consists of *N* sequential blocks, the number of which, *N*, is proportional to the input image size according to *N* = log_2_(imagewidth) – 2 (rounded to the nearest integer). For example, a CNN for size 128 × 128 image inputs would use five blocks. The layers in each block have twice the number of filters as the previous block. The output of the final block is flattened and passed into a dropout layer with keep probability of 0.5. The dropout layer acts to prevent over-fitting of the training data [ref]. Following dropout is a 512-length dense layer with ReLU activation and then the final dense layer with SoftMax activation and the same dimension as the number of classes.

Foraminifera images contain many structural features that are repeated at various locations, differing only by their orientation, such as edges at the particle boundary, lines delineating chambers, corners where chambers meet and so on. This means there is some duplication in the convolution layer filters due to having to account for the same feature at different orientations. We augment the network wit cyclic layers (Dieleman et al., 2016) to reduce this redundancy. A cyclic slice layer is inserted after the image input, creating four parallel paths corresponding to rotations of [0, 90, 180, 270]. After each convolutional block, the output of each path is rotated back, combined, and sliced again (cyclic roll). Then after the first dense layer, the four paths are combined by choosing the maximum value from each path (cyclic pool). In short, this creates for rotations of each filter in the convolutional layers.

Our second design, *ResNet-Cyclic*, has the same topology as the *BaseCyclic* design, except that a ResNet-style skip connection is introduced for each block. The motivation behind the ResNet skip connection is to allow important smaller features to skip ahead to the final dense layers rather then be forced through subsequent convolutional layers.

The main tuneable parameter is the number of filters in convolutional layers of the first block (the number is doubled in subsequent blocks). We typically use either 4, 8 or 16 filters, which because of the cyclic layers, roughly corresponds to 16, 32 or 64 filters without cyclic layers.

### C. Full network training

Full network training involves training a CNN from scratch. Because of the smaller size core-specific foraminifera training sets, choose some of the smaller existing CNNs for use in full network training, in particular ResNet18, SE-ResNet18, MobileNetV2 and DenseNet121, as well as out *Base-Cyclic* and *ResNet-Cyclic* designs.

### D. Transfer Learning

An alternative to full network training is transfer learning, and it has been employed in other approaches to foraminifera classifications with CNNs, such as [REF].

Transfer learning takes an existing CNN that has already been trained on another image set (typically ImageNet [ref]), freezes some or all of the convolutional layers so that their weights can no longer be changed, the re-trains the network on a new training set specific to the task. The idea is that the low-level image features in the earlier stages of the network are common across different classification problems, and therefore it is not necessary to relearn them.

Since the convolutional layers are frozen, our approach to transfer learning is to pre-calculate the vector output of the convolutional layers for each image using global average pooling, train the final dense layers as a separate network, and then recombine them to give the final trained CNN. This method is much faster than training the full network, and is feasible on systems without a GPU. The downside is that random augmentation during training can no longer be used, as this requires the vectors to be recalculated each iteration.

For the trainable dense layers of the network, we use the same configuration as (Zhong et al., 2018; Mitra et al., 2019) consisting of a dropout layer with keep probability 0.05, size 512 dense layer, dropout layer with keep probability 0.15, size 512 dense layer and then a final dense layer with SoftMax activation for the class predictions.

All the CNNs in the Keras library are available with pre-trained weights from ImageNet, and are ideal to use for transfer learning. Each of these networks expects a different scaling and format of the input image, however. For example, ResNet expects BGR images scaled to approximately in the range [−127,127] which DenseNet121 expects RGB images scaled approximately in the range [−0.5, 0.5].

In a production environment we do not wish to have to keep track of which image type and scaling is necessary for which CNN transfer learning topology. Therefore in our implementation we add some extra layers to the beginning of each network to transform the input images (pre-processed and rescaled to [0,1]) to the appropriate format.

For transfer learning we again consider the smaller networks with pre-trained ImageNet weights, specifically VGG19, ResNet50, Xception, InceptionResNet, MobileNetV2, DenseNet121 and NASNetMobile

### E. Cyclic Transfer Tearning

Since our method of transfer learning does not use image augmentation, we also created a variation of the transfer learning topology that includes cyclic layers, so as to intorduce some robustness to foraminifera 2D rotation. In this variation, a cyclic slice layer is inserted before the pre-trained convolutional layers, and a cyclic dense pooling layer is inserted after the global average pooling layer.

### F. Input Dimensions

A final consideration in the topology is the input dimensions of the images fed to the CNN. As foraminifera can appear at any 2D rotation in a slide image, and thus have no dominant orientation, we use a square shaped input.

The choice of input dimensions is limited by the image resolution; using a size greater than the maximum size of the images will require magnification and therefore adds no new information. On the other hand, reducing the input size is useful as it means faster convolutions and thus faster training. This may result in an accuracy penalty if important image features needed to discriminate classes, such as pore texture or secondary apertures, are lost.

Unless colour is a discriminating feature in the image set, we prefer to use single channel (greyscale) images where possible, as it removes colour variations that may adversely affect classification, for example when applying the network to another image set with different colour balance. The transfer learning networks expect three channel colour images, however.

## V. TRAINING

The CNNs are trained using the SoftMax cross-entropy loss function on the predicted labels. We use Adam (adaptive moment) optimisation (Kingma and Ba, 2014) an initial learning rate of 0.001, as it this been found to be good starting point for other image sets (Wilson et al., 2017). A batch size of 64 is used for most CNNs except for larger networks with bigger input image sizes, where it is reduced to 32 or 16 if there are memory constraints.

Training is performed via custom Python scripts and uses the Tensorflow library.

### A. Pre-processing

Three parameter-less pre-processing steps are applied to the images after they are loaded for training. The same settings are laso used for inference.

1. The image intensity is rescaled into the range to [0,1]. That is, an 8-bit image is divided by 255, a 16-bit image by 65535, etc. This removes variance due to bit depth.
2. Any non-square images are padded symmetrically to make them square. A constant padding fill value is used, equal to median value of all the pixels lying on the edge of the image. The edge pixels are used because they are normally background pixels, as the foraminfera particle should be located in the centre of the image.
3. Finally, the square image is resized to the input dimensions of the network, using bilinear interpolation.

### B. Augmentation

When using full network training, augmentation transforms are applied to images during the training stage to increase the robustness of the CNN (e.g., Simard et al., 2003) to image variations. We apply augmentation to simulate some of the variances that arise from the foraminifera imaging system:

1. Random rotation between 0 and 360 degrees.
2. Random gain (brightness) chosen from the set *β* ∈ {0.8, 1.0, 1.2} using the formula 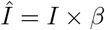
3. Random gamma (contrast) from the set *γ* = {0.5, 1.0, 2.0} using the formula 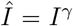. This requires the input images to be in the range [0,1].
4. Random zoom chosen from the set {0.9, 1.0, 1.1}. Values about 1.1 are not use as they would clip the image.

Augmentation is performed in parallel on GPU during training and does not noticeably increase training time.

### C. Class Weighting

The loss function is also weighted inversely according to the count for images in each class. This is to ensure the CNNs are not overfit on the classes with more numerous examples, and boost the accuracy on the more rare foraminifera that may not be very abundant.

The weighting per class is given by the geometric mean of all the class counts, divided by the individual class count. That is,

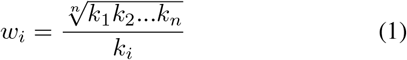

where *w*_*i*_ is the weight for class *i* and *k*_*i*_ is the class count.

### D. Automatic Learning Rate

A periodic decrease in learning rate tends to increase classification accuracy (e.g., He et al., 2016), and the number of batches must be large enough that training completes. We employ an automated method of scheduling learning rate drops and stopping training, based the approach used in the dlib library (King, 2009).

The loss, *y*_*i*_, after each training batch *x*_*i*_ in the last *n* batches, with index *i* ∈ {0‥*n* − 1}, is modelled as a linear function with slope, *m*, and intercept, *c*, corrupted by Gaussian noise, *ϵ*:

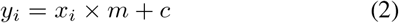

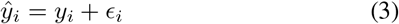

The slope 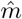 of the noisy loss signal of the last *n* values is a Gaussian random variable with the distribution

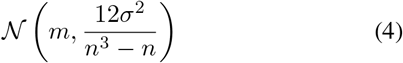

where

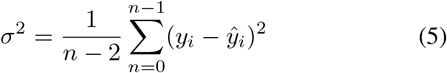

The probability, *P*, of the true slope (*m*) being below 0, that is, the training score is improving, is given by the Gaussian cumulative distribution function

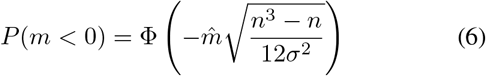

where 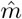 is found using linear regression over 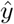.

After each batch, we calculate *P*, and if *P* < 0.51 we assume that training is no longer improving, and so the learning rate is dropped by half. Calculation of *P* is then paused until another *n* batches have been processed. This process is repeated a specified number of times (drops) then once the loss is again no longer improving, training is stopped. We express the number of batches in terms of number of epochs (complete run through the training set), that is,

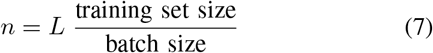

where *L* is the number of epochs.

Is this way, changing the size of a batch does not affect the actual number of images considered when calculating *P*. The number of epochs and drops are tuned to the training set and we find those with large numbers of images per class require fewer epochs. We refer to our automatic learning rate scheduler as ALRS henceforth.

For smaller training sets (¡5000) images with fewer examples per class we use 40 epochs and 4 drops. For larger sets, the number of epochs is dropped, e.g. 10 epochs for ¿10000 images. More drops are added if the noise in the validation accuracy is still significant after four drops.

## VI. TUNING AND EVALUATION

Hyper-parameter tuning is used to obtain the best performing network within the constraints of acceptable training times, inference times and computing requirements.

In our approach we restrict the number of parameters to explore to the following:

- Using pre-calculated vector transfer learning or full network training.
- The network topology, such as type and whether to use cyclic layers.
- The image input size and colour
- The number of filters used in our custom *Base-Cyclic* and *ResNet-Cyclic* CNNs.

To measure the performance of the CNN, a random 20% subset of the training set is reserved for validation, and the remaining 80% is used for training. The validation set is unseen by the CNN during training, and is used to calculate the following measures:

- Overall accuracy: the percentage of images in the validation set that were correctly classified by the CNN. Higher accuracy means better classification performance. We also calculate some per-class measures and report them averaged over all classes:

– Precision: The percentage of images identified into a class that actually belong to the class.
– Recall: The percentage of images in a class that were correctly identified (per-class accuracy)
– f1 score: The average of precision and recall.
- Training time: The time to train the network, including vector calculation in the case of transfer learning. A long training time can reduce the efficiency of the workflow, especially during a hyper-parameter search where multiple trainings are performed. Networks with very short training times may be possible to train on a computer without GPU.
- Inference time: The time to classify a single image. Longer inference time means longer to classify large down-core image sets.

The best performing network is “frozen”, whereby trainable variables are replaced with constants, and saved in protobuf format. An XML file is created with metadata about the network, such as the input size and class names, so that all the information necessary to be able to use the network for classification is present, and thus the CNN can be readily shared for with other users. An example XML file is [here]. The frozen network is now ready for use in classification.

An optional step is to the train the entire image set (both training and validation) on the best performing network. Since there are no validation images the accuracy cannot be measured, however, one would expect that the extra images should improve classification performance on new images.

## VII. CLASSIFICATION

Finally, the chosen trained network is used to classify the larger down-core image set.

Images in the down-core set are arranged into folders by depth. Each is pre-processed as for training (Section V-A), and passed into the CNN to calculate the final layer SoftMax output. The output is a vector of prediction scores, one for each class, ranging from 0 to 1, with all scores adding to 1. We consider a score above a fixed threshold as a positive classification for the class. If no scores are above the threshold, the image is classed as “unsure”. The threshold must be chosen from the range (0.5, 1.0] as then only one class will be above the value. We use a threshold of 0.8. These operations are performed using the *ParticleTrieur* program developed at CEREGE.

## VIII. DEMONSTRATION ON ENDLESS FORAMS

In this section we demonstrate the performance of the different CNN topologies and their parameters, as well as the training method, with a hyper-parameter search using the large, publicly available “endless forams” planktonic foraminifera image set(Hsiang et al., 2019).

The image set consists of 27,729 colour images in 35 species classes, ranging from 4 specimens (Globigerinella adamsi) to 5914 specimens (Globigerinoides ruber) in each class. We excluded five classes, Globigerinella adamsi (4), Globigerinita uvula (7), Tenuitella iota (8), Hastigerina pelagica (13), and Globorotalia ungulata (25) because they had less than 40 images, meaning that only 1 to 8 images are available for validation and thus likely not a reliable measurement.

Each image was pre-processed to remove the white meta-data panel at the bottom of the image and the black border around the particle, so that only the real photographic part of the image remained, then padded to make them square using the method in Section V-A. The processed images are available for download from https://1drv.ms/u/s!AiQM7sVIv7fah4UKCwkj2wMbG2doUA?e=QnbK5I

Both the transfer learning and full network training approaches were investigated using the CNNs described in Section IV. Training was run using Tensoflow 1.14.1 and Python 3.7, on a Windows 10 desktop computer with NVIDIA RTX 2080 Ti GPU, AMD Ryzen 2700X CPU, Sandisk 970 EVO SSD, and 32GB of RAM.

### A. Transfer Learning

To begin, a variety of the transfer learning networks were constructed according to Section IV-D, but without cyclic layers. Colour images at the default size for each network (224 × 224 except for 299 × 299 for Xception and NASNet) were used, and 10 epochs and 4 drops were set for the adaptive learning rate system. Training was repeated five times using 5-fold cross validation, and each performance measure was average across the set.

The ResNet50 network outperformed all others for accuracy (81.8%) and took only 198 seconds to train. MobileNetV2 was the fastest to train, thanks to having the fastest inference time (1.25ms compared to 2.11ms for ResNet50) which makes pre-calculating the vectors faster (Table I).

**TABLE I.** RESULTS OF TRAINING VARIOUS TRANSFER LEARNING NETWORKS WITH THE ENDLESS FORAMS TRAINING SET (COLOUR). TRAINING TIME INCLUDES PRE-CALCULATION OF THE TRANSFER LEARNING VECTORS. *224 × 224 × 3 IMAGES, † 299 × 299 × 3 IMAGES.

| CNN | Acc. | Prec. | Rec. | f1 | Train. time (s) | Epochs | Inf. time (ms) |
| --- | --- | --- | --- | --- | --- | --- | --- |
| resnet50* | **81.8** | **76.7** | **71.4** | **73.4** | 198 | 105 | 2.11 |
| densenet121* | 80.2 | 75.1 | 69.2 | 71.3 | 229 | 119 | 2.34 |
| vgg19* | 77.1 | 70.5 | 64.9 | 66.9 | 187 | 110 | 2.56 |
| xception† | 76.6 | 68.1 | 63.7 | 65.3 | 372 | **102** | 3.91 |
| inceptionresnetV2† | 77.7 | 69.5 | 64.8 | 66.4 | 714 | 149 | 5.83 |
| mobilenetV2* | 77.7 | 70.0 | 65.2 | 66.8 | **166** | 103 | **1.25** |
| nasnetmobile* | 72.3 | 63.8 | 57.6 | 59.6 | 268 | 119 | 2.36 |

Give that ResNet50 had greatest accuracy, we explored the effect of image size and colour, and cyclic layers on this topology (Table II). The transfer learning networks require a three-channel image, so for greyscale images we duplicated the single channel three times. Using greyscale images dropped the accuracy to 79.9%, however, so greyscale was dropped from further consideration.

**TABLE II.** RESULTS OF TRAINING ON THE ENDLESS FORAMS TRAINING SET (COLOUR) USING THE BEST PERFORMING TRANSFER LEARNING NETWORK WITH AND WITHOUT CYCLIC LAYERS, AND FOR DIFFERENT INPUT IMAGE SIZES.

| Image | CNN | Acc. | Prec. | Rec. | f1 | Train. time (s) | Epochs | Inf. time (ms) |
| --- | --- | --- | --- | --- | --- | --- | --- | --- |
| 224 x 224 (grey) | resnet50 | 79.9 | 72.9 | 67.5 | 69.3 | 212 | 103 | 2.05 |
| 64 x 64 | resnet50 | 64.0 | 55.4 | 50.8 | 52.3 | 173 | 124 | **0.32** |
| 128 x 128 | resnet50 | 75.9 | 69.7 | 63.7 | 65.8 | **155** | 103 | 0.74 |
| 224 x 224 | resnet50 | 82.0 | 76.9 | 73.2 | 74.6 | 183 | 99 | 1.95 |
| 320 x 320 | resnet50 | 84.1 | 80.4 | 75.3 | 77.1 | 242 | 97 | 5.80 |
| 416 x 416 | resnet50 | 85.2 | 80.6 | 76.3 | 77.9 | 384 | 109 | 6.48 |
| 64 x 64 | resnet50 cyclic | 71.5 | 65.5 | 60.5 | 62.1 | 201 | 138 | 0.64 |
| 128 x 128 | resnet50 cyclic | 80.6 | 75.2 | 71.0 | 72.6 | 210 | 118 | 1.85 |
| 224 x 224 | resnet50 cyclic | 85.2 | 80.7 | 77.4 | 78.6 | 299 | 111 | 5.47 |
| 320 x 320 | resnet50 cyclic | 86.9 | 83.3 | 80.3 | 81.4 | 498 | 113 | 11.52 |
| 416 x 416 | resnet50 cyclic | 87.2 | 84.2 | 79.8 | 81.5 | 795 | 119 | 18.87 |

Interestingly, increasing the image size improved accuracy, up to 85.2% for 416 × 416 images. However, this significantly increases memory requirements, with the training set (represented as 16 bit floating point numbers) taking approximately 28 GB of memory. The inference time increased with the image area, taking more than three times longer to process 416 × 416 images (6.48ms) compared to 224 × 224 images (1.95ms).

Using cyclic layers improved the accuracy for all images sizes, with a maximum accuracy of 87.2% for 416 × 416 images. The improvement comes at the expense of inference time, which more than doubled.

### B. Full Network Training

We also investigated full network training of the ResNet18 network and our custom base cyclic and ResNet cyclic topologies, for 64 × 64 and 128 × 128 **greyscale** images. For our networks, the number of filters in the first block was varied between 4 and 16 III).

For both 64 × 64 and 128 × 128 images, the base cyclic design with 16 filters gave the best overall accuracy with 87.5% while the the ResNet cyclic with 16 filters gave the best class-averaged precision and recall. In all cases, using 128 × 128 images gave higher accuracy, up to 90.3%.

Training time increased proportionally with image size and number of filters, and the *ResNet-Cyclic* network took longer than the base cyclic network, up to almost 3.5 hours for one configuration. The long training time is why larger image sizes were not investigated further for this dataset, however, it may be feasible for smaller sets of just a few thousand images. Inference time was low for all configurations, ranging from 0.09 ms to 0.68 ms, except for the ResNet18 network with was 2.05 ms.

### C. Comparison

The full network training gave better accuracy than the transfer learning methods at the expense of much longer training times. The likely due to the use of augmentation with the full network training. Using augmentation with transfer learning would require calculation of the vector output every time a training image is fed into the network, rather than a single pre-calculation at the start. Since the inference times of the transfer learning networks were around 2 to 10 times longer than longest for the *Base-Cyclic* and *ResNet-Cyclic* full networks, we can infer that using augmentation with transfer learning would result in training times much longer than the already long times for the full networks.

The short training time for the transfer learning networks is advantageous for quickly investigating the expected accuracy for a particular dataset. In particular, looking at the individual class accuracies can help identify any deficiencies in the training set, for example, a low accuracy suggests more images are needed for a particular class. Furthermore, a short training time indicates low computation complexity, and we find that the transfer learning approach is feasible on computers without a high-performance GPU.

The long training time of full network training of the *Base-Cyclic* and *ResNet-Cyclic* networks is offset by their short inference times, which in turn affects how quickly a large down-core image set can be classified. If a trained CNN is shared with other users who wish to classify images on their personal systems with high-performance GPUs, the inference time will be much longer. In this case the long training time is justified for obtaining a shorter inference time.

## IX. APPLICATION TO BENTHIC FORAMINIFERA DATASET (CORE MD02-2508)

The second application was to create a high-resolution analysis of the Holocene interval within Core MD02-2508, retrieved from the North Eastern Pacific during the R/V Marion-Dufresne MD126 MONA (Image VII) campaign in 2002 (Beaufort, 2002).

### A. Down-core Set

A large down-core image set (73544 images) was acquired for core MD02-2508 using the imaging system described in Section II. Some images were taken with the particles on a micropaleontological slide, and some were taken using the MISO particle sorting machine at CEREGE. Individual particles were segmented from these larger images as per our method.

Images were taken of 41 samples from 40cm to 642cm deep. These samples were chosen to cover the Holocene and transition from the last Glacial to Interglacial state (0 - 16000 years ago) as given by the age model for this core. Manual counting of benthic species in this core had already been performed for 37 samples in this depth range (Tetard et al., 2017), then used for comparison.

**TABLE III.** RESULTS OF FULL NETWORK TRAINING ON THE ENDLESS FORAMS TRAINING SET (GREYSCALE) FOR DIFFERENT INPUT SIZES AND NUMBER OF FILTERS.

| Image | CNN | Acc. | Prec. | Rec. | f1 | Train. time (s) | Epochs | Inf. time (ms) |
| --- | --- | --- | --- | --- | --- | --- | --- | --- |
| 64 | resnet18 | 84.9 | 78.0 | 71.8 | 73.6 | 2656 | 229 | 0.15 |
| 64 | base cyclic 4 | 83.1 | 77.3 | 68.6 | 71.0 | **1691** | 213 | **0.09** |
| 64 | resnet cyclic 4 | 83.5 | 77.4 | 70.0 | 72.2 | 2428 | 229 | 0.12 |
| 64 | base cyclic 8 | 86.3 | 80.7 | 73.3 | 75.4 | 2361 | 222 | 0.13 |
| 64 | resnet cyclic 8 | 86.6 | 81.3 | 74.0 | 76.3 | 3888 | 241 | 0.18 |
| 64 | base cyclic 16 | **87.5** | 78.9 | 73.6 | 75.4 | 3173 | 199 | 0.19 |
| 64 | resnet cyclic 16 | 86.9 | **82.3** | **74.1** | **76.5** | 4986 | 191 | 0.28 |
| 128 | resnet18 | 88.5 | 84.1 | 77.8 | 79.9 | **3903** | 198 | 0.30 |
| 128 | base cyclic 4 | 89.1 | 83.6 | 78.4 | 80.1 | 4766 | 214 | **0.26** |
| 128 | resnet cyclic 4 | 87.8 | 82.8 | 76.4 | 78.1 | 7318 | 241 | 0.33 |
| 128 | base cyclic 8 | 90.1 | 83.7 | **79.0** | 80.7 | 5687 | 169 | 0.40 |
| 128 | resnet cyclic 8 | 89.5 | **85.3** | **79.0** | **80.9** | 12495 | 245 | 0.55 |
| 128 | base cyclic 16 | **90.3** | 84.9 | 78.4 | 80.5 | 8693 | 151 | 0.68 |
| 128 | resnet cyclic 16 | 90.1 | 85.1 | 78.7 | 80.8 | 32619 | 214 | 1.63 |

### B. Training Set

A training set was constructed from 15274 images of foraminifera from seven representative samples from cores MD02-2508 and MD02-2519. The images from MD02-2519 (not the core of interest) were used as the core is from a similar location to MD02-2508 and contained a very similar benthic foraminiferal fauna, and the images had already been acquired.

The training images were manually labelled using the *ParticleTrieur* software into 12 benthic species, (main species according to Tetard et al. (2017)), and an “other-benthic” class (grouping the less abundant benthic species), a single catch-all planktonic class, a radiolarian class, and then some non-foraminifera classes such as double (specimens in contact with each other) and fragments (Figure 3). Images ranged from 188 × 188 to 1502 × 1502 pixels in size, corresponding to particles 0.14 mm to 1.1 mm in diameter.

**Fig. 3.**
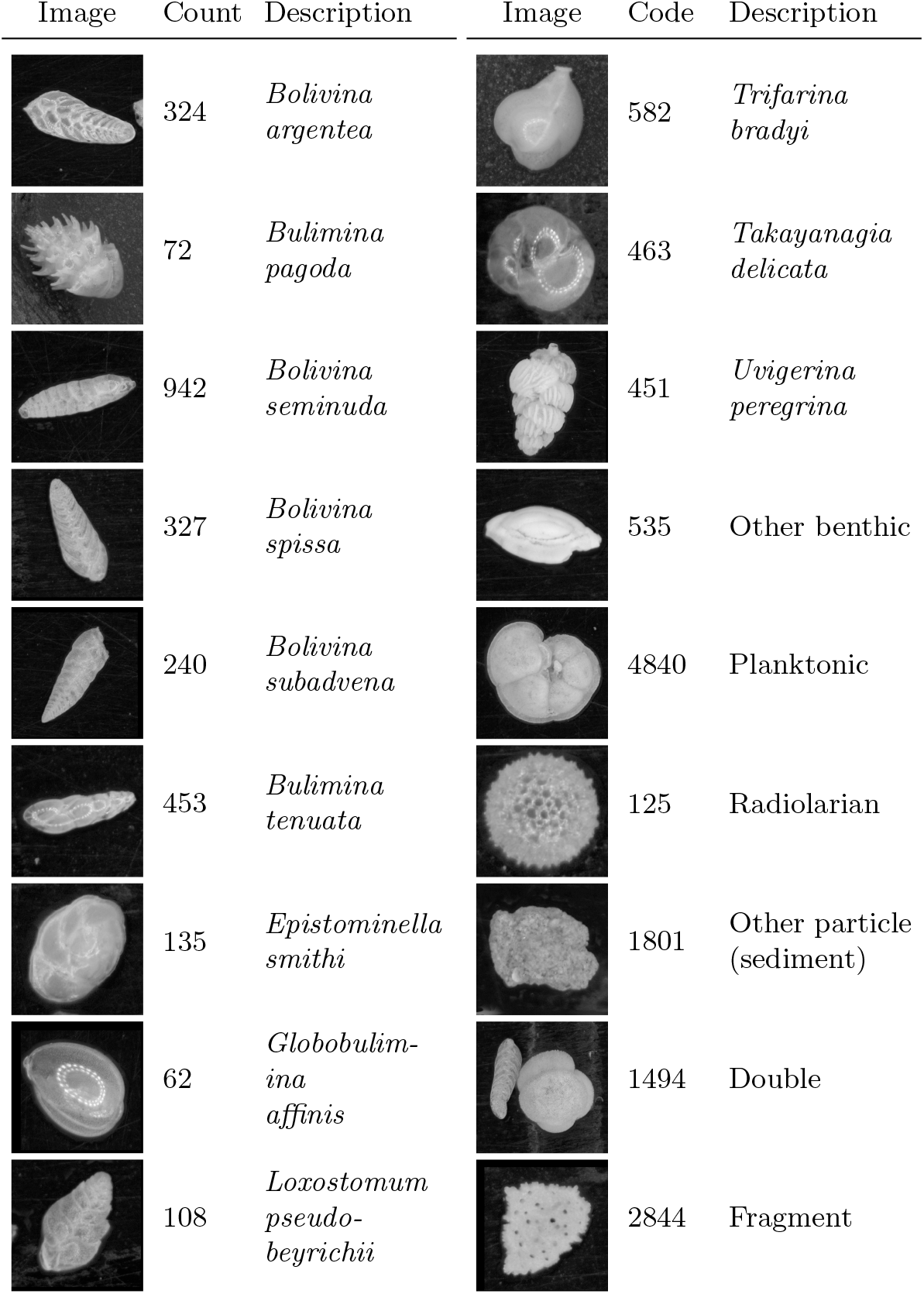
Example image from each class of training set constructed from cores MD02-2508 and MD02-2519, classified mainly by species. There are 15274 images in total.

### C. Classification

A small search was conducted to find the best performing CNN, using an 80% train / 20% validation split, 10 ALR epochs and either the ResNet50 transfer learning with and without cyclic layers, and the Base Cyclic network with 4 or 8 filters.

The Base Cyclic network with 8 filters gave the best accuracy (89%) with most classes having above 75%. There was some confusion between similar looking *Bolivina* benthic species, *B. spissa*, *B. subadvena* and *B. seminuda*. Precision and recall (per-class accuracy) tended to be higher for those classes with a high count in the training set. Almost all classes had some confusion with the fragment class (Figure 4).

**Fig. 4.**
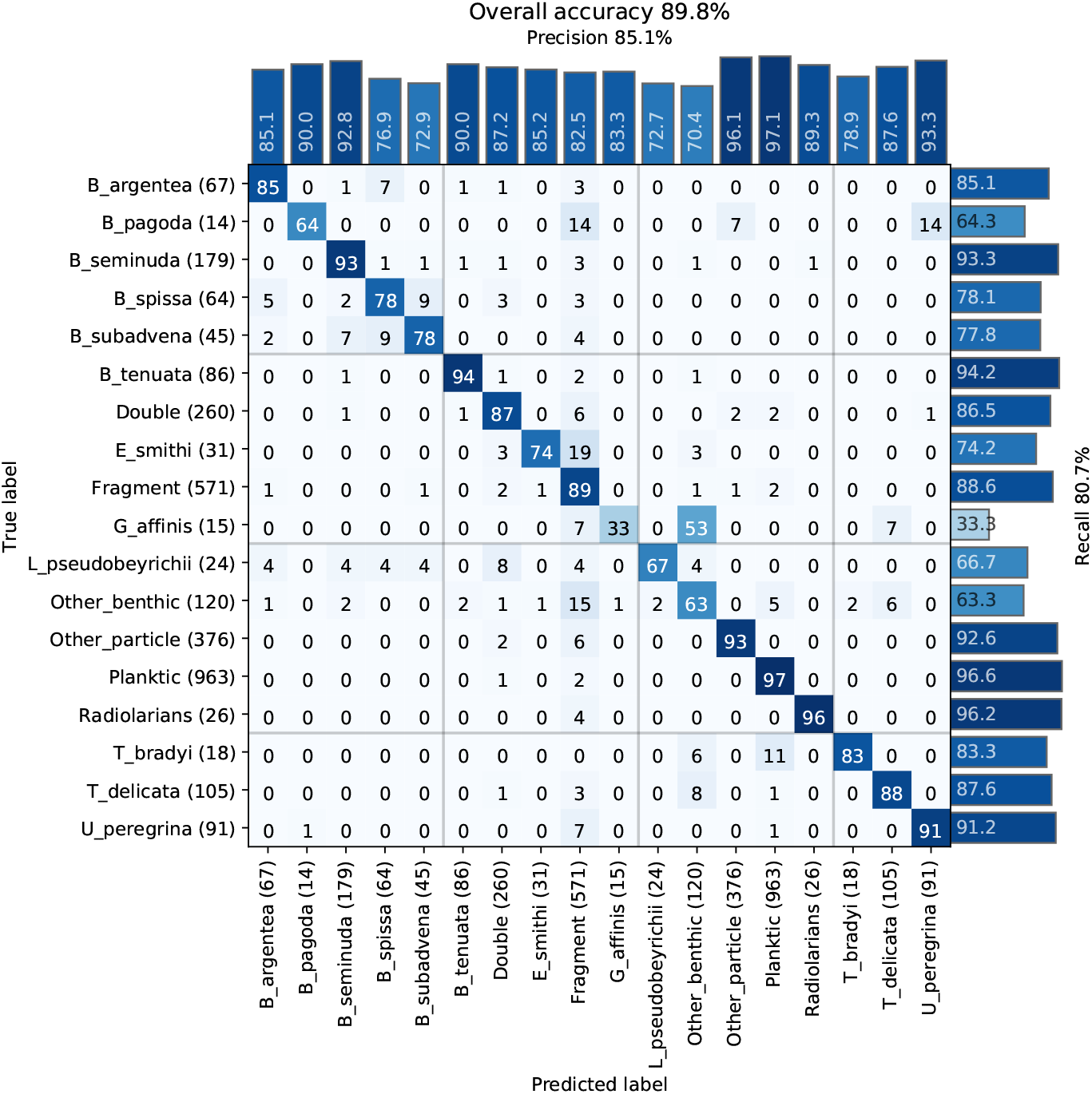
Confusion matrix of one training run of the core MD02-2508 training set showing the percentage of images in each class (rows) classified into a particular class (columns). The number of images in the validation set for each class is shown in brackets next to the class label.

A review of the training set found errors such as mis-labelling and duplicate images (due to a slight overlap in the images acquired using an automated stage) that were labelled into different classes, and these may have negatively affected the accuracy. Furthermore, the presence of plastic core liner or sediment particles touching the foraminifera of interest occasionally resulted in the image being classified into either the “double” class, or another class with similar shape to their combined appearance. Likewise, the variability of fragmentation from slight damage to a single chamber to larger damage affecting a number of chambers may explain why some images in each class were classified as fragments (Figure 5).

**Fig. 5.**
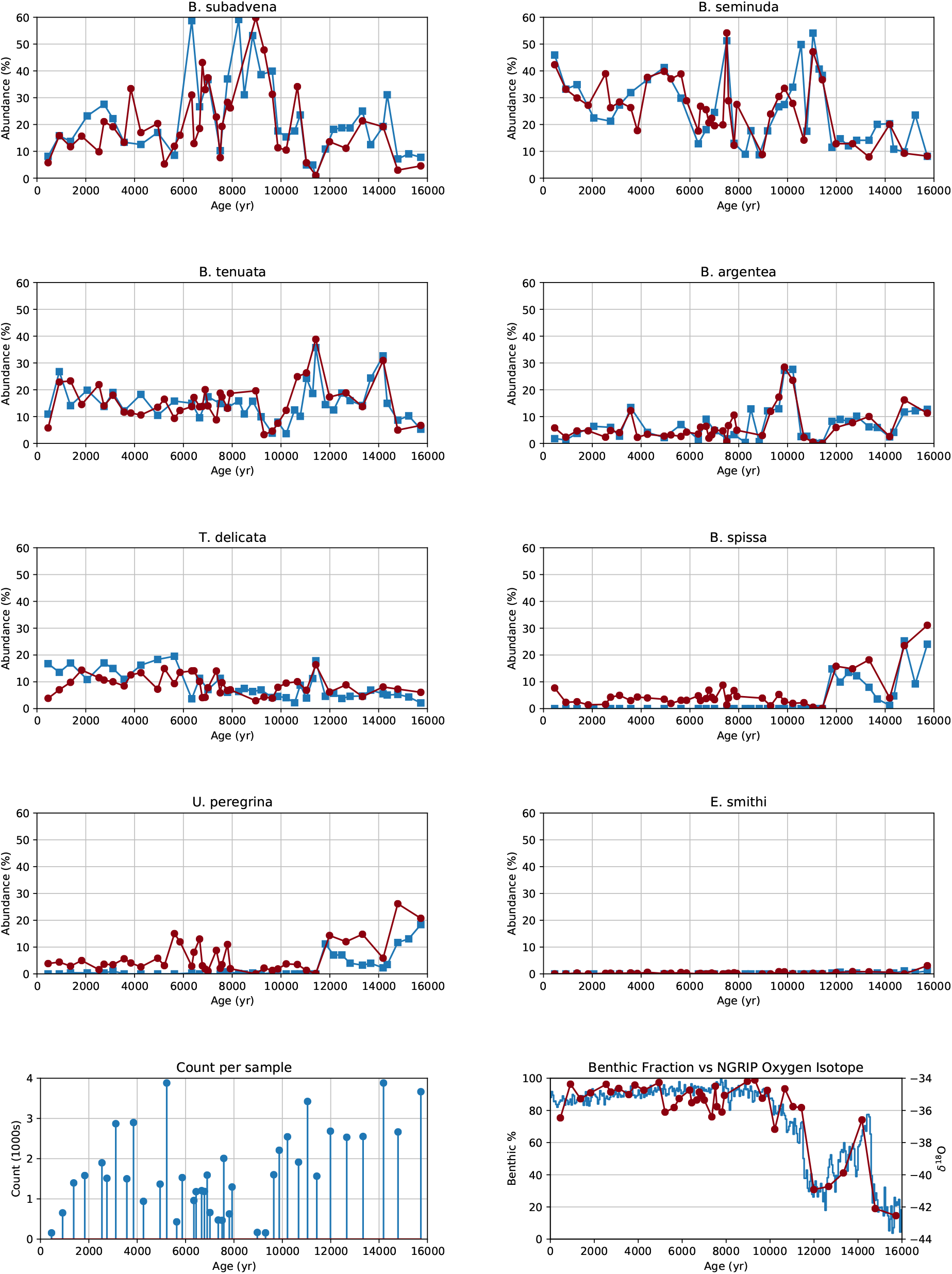
Relative abundance of eight benthic species in core MD02-2508 (top), image counts per sample (bottom-left) and the benthic foraminifera to whole foraminifera ratio compared to the Greenland oxygen isotopic record (bottom-right).

### D. Abundance

Manual counting of samples from MD02-2508 had previously been performed for every benthic species recovered from this core. For the results of both manual and CNN counting, we separated out eight of the main species that are of interest for paleo-oceanographic reconstructions (Tetard et al., 2017), and placed the rest into an “other benthic” class. The relative abundance was then calculated for each benthic group compared to the total benthic count, for each method.

Since the planktonic classes were undifferentiated and no manual counting had been performed for them, we instead calculated the percentage of benthic foraminifera to whole foraminifera (planktonic + benthic) from the CNN counts over the same period, and compared the shape of this signal to the Greenland oxygen isotopic record.

The signals obtained using CNN classification had similar dynamic characteristics to those from manual counting (Figure 5).

- Counts for *B. argentea* are higher than for the human counts in the more recent samples, however the signal exhibits the same dynamics, with the same peaks in abundance around 10000, 3500, and 1200 yrs BP.
- *Bolivina seminuda* also shows similar abundance with the human counts, with gradually increasing abundance towards the modern day. The short spike in abundance around 7500 yrs BP is also visible.
- Human counting of *B. spissa* is zero for the Holocene, and this species is usually absent from this area during this period. The CNN finds some 10% of specimens as *B. spissa*, suggesting all of these images may have been misclassified
- Counts of *B. subadvena* show similar absolute abundance and dynamics as human counting, except for a peak around 6500 ya.
- *Buliminella tenuata* shows the same large peaks at 11500 and 14000 yrs BP. The transition from almost zero abundance at 9500 yrs BP to above 30% abundance at 11500 yrs BP is also much smoother for the CNN derived results, suggesting that the larger number of images processed result in less noise for the highly abundant species.
- Human counts for *E. smithi* during the Holocene are zero, suggesting the absence of this species during this time in this particular area. The CNN counts are also low during this period, in contrast to the *B. spissa* signal.
- Similarly, the non-zero CNN counts of *U. peregrina* during the Holocene are likely due to the presence of a morphologically close species: *U. striata*.
- The CNN signal for *T. delicata* also follows the human counting, with a peak at 11500 yrs BP present in both results.
- Both human and CNN abundances show other benthic species at around 10% during the Holocene and 20% before it. The CNN counts are much smoother.
- CNN counts show that the percentage of benthic foraminifera was very high during the Holocene, dropping off around 11500 ya. The dynamics of the signal very closely match that of the Greenland oxygen isotopic record, correlating with other studies that show that benthic foraminifera abundance and marine productivity were higher during warm periods, especially the Holocene, in this area (Cartapanis et al., 2011, 2014, Tetard et al, 2017). The results also indicate the CNN can successfully discriminate benthic from planktonic species, likely due to the greater difference in morphology compared to within benthic species.

### E. Discussion

Applied to the down core images, the dynamics of each abundance signal was similar between the CNN and manual counting. However, we notice that the strongest bias in most species is likely caused by false positives. We inspected the misclassified images to try to find the source of the errors. As with the training results, the various species of the *Bolivina* genus were generally confused with each other. In particular, many specimens of B. subadvena were misclassified as *B. spissa*, causing the non-zero counts for this species during the Holocene.

One possible explanation is that the intraspecific morphometric variability for species *B. argentea*, *B. spissa* and *B. subadvena* can be higher than the interspecific variability between these species. For example, the microspheric forms of different species can appear more similar than with the microspheric and macrospheric forms of the same species. As a consequence, the identification and discrimination of these forms can be difficult even for a taxonomist, and the CNN also mistakes these species more often. This can also explain why *E. smithi*, which was also not present in the core during the Holocene, did not have a strong false positive bias, as few other classes were confused with it during training.

We note some species can be discriminated under a stereoscopic microscope by their flatness, (e.g. *B. spissa* and *B. argentea* versus *B. subadvena* and *B. seminuda*), which helps for manual identification, but this depth information is lost for the 2D images in our automated approach. This bias is thus largely dataset dependant as out of the 8 main species analysed in this study, four belong to the same genus (i.e. *Bolivina*). Thus the dataset provides a good case study on the perfomance of a CNN classifier; overall the most of the classes were correctly identified.

## X. CONCLUSION

In this paper we have present a method for analysing downcore foraminifera image set using deep convolutional neural networks. The performance of transfer learning and full network training for publicly available CNNs, as well as our custom *Base-Cyclic* and *ResNet-Cyclic* designs, were demonstrated on the Endless Forams image set, as well as our core-specific benthic training set. The transfer learning approach is fast to train and gives good accuracy without augmentation. Full network training is much slower to train, but our *Base-Cyclic* design gives as good or better accuracy with faster inference time. Our approach has been to use transfer learning when a quick CNN is needed to aide with manual labelling, and full network training to create the final network used for down-core analysis.

This method is currently used at the CEREGE laboratory, and can also be applied to classify other image types, such as pollen or plankton. An important observation we have made during this time is that even with heavy image augmentation, classifying images using a CNN trained on images from a different imaging system is not as accurate as classifing with those obtained from the same system. In particular, a change in background can cause gross misclassification, e.g. a particle imaged on a micropaleontogical tray compare to one imaged in our MISO foraminifera sorting machine. We recommend keeping the same imaging settings for both the down-core and training image sets.

Likewise, one should optimize the training set according to the sediment or core under analysis. This is important in three ways: the training set should (i) incorporate all the main taxa and their morphological variants, (ii) have undergone the same early diagenetic history, to ensure that the range of dissolution, early pyritization (which can affect structure), colour, and translucency are included in the morphological variability, and (iii) include non-foraminifera artefacts that could affect classification, such as particles (e.g. plastic core liner or sediment) or specifics of the acquisition system (e.g. ring light pattern).

The example application shows that very large throughput is possible with an automated system. A few hours labelling a well-constructed training set can save months of time manually counting specimens. Furthermore, the CNN obtained can be re-purposed to aide in constructing other training sets by using the predictions to suggest labels.

The software program *ParticleTrieur* which was used to manually label the training set and automatically classify the down-core set is available with tutorial from http://particle-classification.readthedocs.io, and the python scripts to train the CNNs in this paper are available from http://www.github.com/microfossil/particle-classification and can be install using the *pip* Python package.

## XI. FUTURE WORK

One major pitfall of the CNN classification is the requirement of a uniform size input. Size information of foraminifera is lost in this process, though this is might be important for some morpho-classes which can be discriminated by size.

Other morphometric information that is not well-represented by a CNN could also assist in classification, for example, chamber count and texture distribution. Although this requires feature engineering rather than learning, the measurements are interpretable and thus relevant to taxonomists and rule-based classification. This is opposed to CNN features which are local, generally not interpretable, and not necessarily consistent between image sets.

Likewise, specimen thickness could help discriminate round and flat species, such as *B. spissa* and *B. subadvena* in the benthic image set. Thickness can be estimated from the depth map calculated when performing multi-focal image fusion (needs ref). In future work we propose to combine morphometric features, depth mapping and a CNN in a new classification system.

## CONTRIBUTIONS

R. Marchant developed the system, performed the experiments, acquired images for core MD02-2508 and was the primary author of the paper, M. Tetard acquired and expertly labelled the images for core MD02-2508 and edited the paper, A. Pratawi acquired and labelled the images for core MD02-2508B, and T. de Garidel-Thoron organised the project and its funding and wrote the paper.

## ACKNOWLEDGEMENTS

This work was funded under a Marie Curie PRESTIGE grant and the ANR PRCE FIRST project.

## REFERENCES

N. Barbarin. La reconnaissance automatisée des nannofossiles calcaires du cénozoïque. PhD thesis, 2014.

L. Beaufort. MD 126 / MONA cruise, RV Marion Dufresne, 2002.

L. Beaufort and D. Dollfus. Automatic recognition of coccoliths by dynamical neural networks. Marine Micropaleontology, 51(1–2):57–73, apr 2004. ISSN 03778398. doi: 10.1016/j.marmicro.2003.09.003. URL http://linkinghub.elsevier.com/retrieve/pii/S0377839803001038.

J. Bollmann, P. S. Quinn, M. Vela, B. Brabec, S. Brechner, M. Y. Cortés, H. Hilbrecht, D. N. Schmidt, R. Schiebel, and H. R. Thierstein. Image Analysis, Sediments and Paleoenvironments, volume 7 of Developments in Paleoenvironmental Research. Kluwer Academic Publishers, Dordrecht, 2005. ISBN 1-4020-2061-9. doi: 10.1007/1-4020-2122-4. URL http://link.springer.com/10.1007/1-4020-2122-4.

P. Culverhouse, R. Simpson, R. Ellis, J. Lindley, R. Williams, T. Parisini, B. Reguera, I. Bravo, R. Zoppoli, G. Earnshaw, H. McCall, and G. Smith. Automatic classification of field-collected dinoflagellates by artificial neural network. Marine Ecology Progress Series, 139:281–287, 1996. doi: 10.3354/meps139307.

P. Culverhouse, R. Williams, B. Reguera, V. Herry, and S. Gonz?lez-Gil. Do experts make mistakes? A comparison of human and machine identification of dinoflagellates. Marine Ecology Progress Series, 247:17–25, 2003. ISSN 0171-8630. doi: 10.3354/meps247017. URL http://www.int-res.com/abstracts/meps/v247/p17-25/.

S. Dieleman, J. De Fauw, and K. Kavukcuoglu. Exploiting Cyclic Symmetry in Convolutional Neural Networks. Arxiv, page 10, feb 2016. ISSN 1938-7228. URL http://arxiv.org/abs/1602.02660.

D. Dollfus and L. Beaufort. Fat neural network for recognition of position-normalised objects. Neural Networks, 12(3):553–560, 1999. ISSN 08936080. doi: 10.1016/S0893-6080(99)00011-8.

K. He, X. Zhang, S. Ren, and J. Sun. Deep Residual Learning for Image Recognition. Multimedia Tools and Applications, pages 1–17, dec 2015. ISSN 1380-7501. doi: 10.1007/s11042-017-4440-4. URL http://arxiv.org/abs/1512.03385 http://link.springer.com/10.1007/s11042-017-4440-4.

K. He, X. Zhang, S. Ren, and J. Sun. Identity Mappings in Deep Residual Networks. pages 630–645. 2016. ISBN 9783319464930. doi: 10.1007/978-3-319-46493-0. URL http://link.springer.com/10.1007/978-3-319-46493-0.

D. Hibbett. Evolution Automated Taxon Identification in Systematics: Theory, Approaches and Applications. The Systematics Association Special Volumes Series, Volume 74. Edited by NormanMacLeod. CRC Press. Boca Raton (Florida): Taylor & Francis Group. $99.95. xvii 33. The Quarterly Review of Biology, 84(3):295–296, sep 2009. ISSN 0033-5770. doi: 10.1086/644681. URL http://books.google.com/books?id=Fz6v8YPyQQgC{&}pgis=1 http://www.journals.uchicago.edu/doi/10.1086/644681.

G. E. Hinton, N. Srivastava, A. Krizhevsky, I. Sutskever, and R. R. Salakhutdinov. Improving neural networks by preventing co-adaptation of feature detectors. ArXiv e-prints, pages 1–18, 2012. ISSN 9781467394673. doi: arXiv:1207.0580. URL http://arxiv.org/abs/1207.0580.

A. Y. Hsiang, A. Brombacher, M. C. Rillo, M. J. Mleneck-Vautravers, S. Conn, S. Lordsmith, A. Jentzen, M. J. Henehan, B. Metcalfe, I. S. Fenton, B. S. Wade, L. Fox, J. Meilland, C. V. Davis, U. Baranowski, J. Groeneveld, K. M. Edgar, A. Movellan, T. Aze, H. J. Dowsett, C. G. Miller, N. Rios, and P. M. Hull. Endless Forams: >34,000 Modern Planktonic Foraminiferal Images for Taxonomic Training and Automated Species Recognition Using Convolutional Neural Networks. Paleoceanography and Paleoclimatology, 34(7):1157–1177, 2019. ISSN 2572-4517. doi: 10.1029/2019pa003612.

G. Huang, Z. Liu, and K. Q. Weinberger. Densely Connected Convolutional Networks. arXiv preprint, pages 1–12, 2016. ISSN 0002-9645. doi: 10.1109/CVPR.2017.243. URL http://arxiv.org/abs/1608.06993.

D. E. King. Dlibml: A Machine Learning Toolkit. Journal of Machine Learning Research, 10:17551758, 2009. ISSN 15324435. doi: 10.1145/1577069.1755843.

D. P. Kingma and J. Ba. Adam: A Method for Stochastic Optimization. pages 1–15, 2014. ISSN 09252312. doi: http://doi.acm.org.ezproxy.lib.ucf.edu/10.1145/1830483.1830503. URL http://arxiv.org/abs/1412.6980.

A. Krizhevsky, I. Sutskever, and G. E. Hinton. ImageNet Classification with Deep Convolutional Neural Networks. Advances In Neural Information Processing Systems, pages 1–9, 2012. ISSN 10495258. doi: http://dx.doi.org/10.1016/j.protcy.2014.09.007.

S. Liu, M. Thonnat, and M. Berthod. Automatic classification of planktonic foraminifera by a knowledge-based system. Artificial Intelligence for Applications, 1994., Proceedings of the Tenth Conference on, pages 358–364, 1994.

R. Mitra, T. M. Marchitto, Q. Ge, B. Zhong, B. Kanakiya, M. S. Cook, J. S. Fehrenbacher, J. D. Ortiz, A. Tripati, and E. Lobaton. Automated species-level identification of planktic foraminifera using convolutional neural networks, with comparison to human performance. Marine Micropaleontology, 147:16–24, 2019. ISSN 03778398. doi: 10.1016/j.marmicro.2019.01.005.

O. Russakovsky, J. Deng, H. Su, J. Krause, S. Satheesh, S. Ma, Z. Huang, A. Karpathy, A. Khosla, M. Bernstein, A. C. Berg, and L. Fei-Fei. ImageNet Large Scale Visual Recognition Challenge. International Journal of Computer Vision, 115(3):211–252, 2015. ISSN 15731405. doi: 10.1007/s11263-015-0816-y. URL http://dx.doi.org/10.1007/s11263-015-0816-y.

J. Schmidhuber. Deep Learning in Neural Networks: An Overview. Arxiv, pages 1–88, oct 2014. ISSN 09574174. doi: 10.1016/j.neunet.2014.09.003. URL http://arxiv.org/abs/1404.7828 http://dx.doi.org/10.1016/j.neunet.2014.09.003.

K. Schulze, U. M. Tillich, T. Dandekar, and M. Frohme. PlanktoVision – an automated analysis system for the identification of phytoplankton. BMC Bioinformatics, 14(1):115, 2013. ISSN 1471-2105. doi: 10.1186/1471-2105-14-115. URL http://bmcbioinformatics.biomedcentral.com/articles/10.1186/1471-2105-14-115.

P. Simard, D. Steinkraus, and J. Platt. Best practices for convolutional neural networks applied to visual document analysis. Seventh International Conference on Document Analysis and Recognition, 2003. Proceedings., 1(Icdar):958–963, 2003. ISSN 15205363. doi: 10.1109/ICDAR.2003.1227801. URL http://ieeexplore.ieee.org/lpdocs/epic03/wrapper.htm?arnumber=1227801.

K. Simonyan and A. Zisserman. Very Deep Convolutional Networks for Large-Scale Image Recognition. pages 1–14, sep 2014. ISSN 09505849. doi: 10.1016/j.infsof.2008.09.005. URL http://arxiv.org/abs/1409.1556.

R. Simpson, R. Williams, R. Ellis, and P. Culverhouse. Biological pattern recognition by neural networks. Marine Ecology Progress Series, 79:303–308, 1992.

C. Szegedy, W. Liu, Y. Jia, P. Sermanet, S. Reed, D. Anguelov, D. Erhan, V. Vanhoucke, and A. Rabinovich. Going deeper with convolutions. Proceedings of the IEEE Computer Society Conference on Computer Vision and Pattern Recognition, 07-12-June:1–9, 2015a. ISSN 10636919. doi: 10.1109/CVPR.2015.7298594.

C. Szegedy, V. Vanhoucke, S. Ioffe, J. Shlens, and Z. Wojna. Rethinking the Inception Architecture for Computer Vision. arXiv preprint, 2015b. ISSN 08866236. doi: 10.1002/2014GB005021. URL http://arxiv.org/abs/1512.00567.

M. Tetard, L. Licari, and L. Beaufort. Oxygen history off Baja California over the last 80 kyr: A new foraminiferal-based record. Paleoceanography, 32(3):246–264, 2017. ISSN 19449186. doi: 10.1002/2016PA003034.

A. C. Wilson, R. Roelofs, M. Stern, N. Srebro, and B. Recht. The Marginal Value of Adaptive Gradient Methods in Machine Learning. pages 1–14, 2017. URL http://arxiv.org/abs/1705.08292.

S. Xie, R. Girshick, P. Dollár, Z. Tu, and K. He. Aggregated Residual Transformations for Deep Neural Networks. 2016. URL http://arxiv.org/abs/1611.05431.

S. Yu, P. Saint-Marc, M. Thonnat, and M. Berthod. Feasibility study of automatic identification of planktic foraminifera by computer vision. The Journal of Foraminiferal Research, 26(2):113–123, apr 1996. ISSN 0096-1191. doi: 10.2113/gsjfr.26.2.113. URL http://jfr.geoscienceworld.org/cgi/doi/10.2113/gsjfr.26.2.113.

S. Zagoruyko and N. Komodakis. Wide Residual Networks. Arxiv, 2016. URL http://arxiv.org/abs/1605.07146.

B. Zhong, Q. Ge, B. Kanakiya, R. Mitra, R. M. T. Marchitto, and E. Lobaton. A comparative study of image classification algorithms for Foraminifera identification. 2017 IEEE Symposium Series on Computational Intelligence, SSCI 2017 - Proceedings, 2018-Janua:1–8, 2018. doi: 10.1109/SSCI.2017.8285164.

